# Dynamic protein deacetylation is a limited carbon source for acetyl-CoA-dependent metabolism

**DOI:** 10.1101/2022.12.28.522149

**Authors:** Ioana Soaita, Emily Megill, Daniel Kantner, Adam Chatoff, Zoltan Arany, Nathaniel W Snyder, Kathryn E Wellen, Sophie Trefely

## Abstract

The ability of cells to store and rapidly mobilize energy reserves in response to nutrient availability is essential for survival. Breakdown of carbon stores produces acetyl-coenzyme-A (acetyl-CoA), which fuels various metabolic pathways and is also the acyl donor for protein lysine acetylation. Notably, histone acetylation is sensitive to acetyl-CoA availability and nutrient replete conditions induce a substantial accumulation of acetylation on histones. Deacetylation releases acetate, which can be recycled to acetyl-CoA, suggesting that deacetylation could be mobilized as an acetyl-CoA source to feed downstream metabolic processes under nutrient depletion. While the notion of histones as a metabolic reservoir has been frequently proposed, experimental evidence has been lacking. Therefore, to test this concept directly, we developed an experimental system to trace deacetylation-derived acetate and its incorporation into acetyl-CoA, using ^13^C_2_-acetate in ATP citrate lyase-deficient fibroblasts (*Acly*^*-/-*^ MEFs), which are primarily dependent on acetate for protein acetylation. We find that dynamic protein deacetylation in *Acly*^*-/-*^ MEFs contributes carbons to acetyl-CoA and proximal downstream metabolites. However, there is no significant effect on acyl-CoA pool sizes, and even at maximal acetylation, deacetylation transiently supplies approximately 9% of cellular acetyl-CoA. Together, our data reveal that although protein acetylation is dynamic and sensitive to nutrient availability, its potential for maintaining cellular acetyl-CoA-dependent metabolic pathways is limited compared to cellular demand.

## Introduction

Cells stockpile energy in intracellular stores such as glycogen and lipid droplets for survival across external variations in nutrient availability. Under nutrient deprivation, mobilization of intracellular energy stores generates acetyl-coenzyme-A (acetyl-CoA), an essential metabolite in anabolic and catabolic pathways required for survival. Acetyl-CoA is also the acyl donor for protein acetylation, thus directly influencing protein signaling and epigenetic regulation [1,2]. Notably, bulk histone acetylation is sensitive to acetyl-CoA availability, and the supply of acetyl-CoA for acetylation of specific histone acetylation marks is important for gene regulation in contexts such as cell differentiation [3–10]. However, whether histone acetylation plays roles beyond gene regulation remains poorly understood [11–16].

Homeostatic turnover of histone acetylation occurs over a timeframe of minutes to hours in mammalian cell culture under stable nutrient conditions. This dynamic turnover is facilitated by removal and conversion of acetylation marks to free acetate through the action of lysine deacetylases (KDACs) [17–20]. Free acetate can then be converted to acetyl-CoA by cytosolic/nuclear localized acyl-CoA synthetase short chain family member 2 (ACSS2) and recycled for the acetylation of specific histone marks [13,21–23]. The contribution of protein deacetylation to whole-cell acetyl-CoA and acetyl-CoA-dependent metabolism, however, is unknown. The potential for bulk acetyl sequestration under nutrient replete conditions has led to speculation that protein acetylation may serve as an acetate reservoir that can be mobilized as a metabolic source beyond the nucleus [11,12,14,15].

To test the capacity for protein deacetylation to directly contribute to cellular acetyl-CoA, we developed an experimental system to detect and trace deacetylation-derived acetate incorporation into acetyl-CoA. We employed ^13^C_2_-acetate in mouse embryonic fibroblasts deficient in ATP citrate lyase (*Acly*^*-/-*^ MEFs), which are primarily dependent on acetate for protein acetylation [19]. We measured the maximal contribution of protein-derived acetate carbon units to acetyl-CoA by inducing deacetylation after maximizing protein acetylation. We found that deacetylation transiently supplied approximately 9% of cellular acetyl-CoA. Our results demonstrate that protein deacetylation can directly contribute to acetyl-CoA, but is not acutely a major source of total cellular acetyl-CoA. The experimental system also provides a framework for testing this phenomenon under other physiological or pathological conditions.

## Results

### Estimating the acetate storage capacity of histones

We first sought to estimate the acetate storage capacity of histones under physiological conditions. We focused on histone N-terminal lysine sites, since histone acetylation occurs primarily on lysine residues of disordered N-terminal tail regions [24,25]. We multiplied the number of N-terminal lysine residues (H2A, H2B, H3 and H4) by estimates for the total number of each histone protein per human diploid cell, which gave approximately 3.03x10^−3^ pmol of acetate per cell (Fig 1A; see S1 Table for calculations) [26–28]. We calculated a similar amount of acetate can be stored on histones in a mouse diploid cell (S1 Table) [29]. Total cellular acetyl-CoA has been measured in the range of 2x10^−5^ to 2x10^−4^ pmol/cell (Figs 1 and 2) [4,30]. Therefore, at complete maximum acetylation capacity, we estimate that histone tails have the potential to provide 10-100 times the whole-cell acetyl-CoA, agreeing with previous estimates [12,15]. However, measurements (relative and stoichiometric) of acetylation occupancy at specific sites indicate that physiological occupancy averages between 4-13% under standard culture conditions across primary and immortalized cell lines (Fig 1A; S1 Table) [17,30–36]. We therefore estimate that 1x10^−4^ to 4x10^−4^ pmol acetyl-CoA per cell (∼0.5-20 acetyl-CoA pool sizes) could be stored on histone tails under standard culture conditions. Histones likely account for at least half the protein bound acetate in the cell [34,37–39] (S2 Table), so even partial deacetylation of histone and non-histone proteins at physiological acetylation levels could potentially serve as a significant source of acetate and acetyl-CoA. Importantly, the fast turnover of acetyl-CoA (half-life <5 minutes) and rapid flux into downstream metabolites [40] suggests that deacetylation is not likely to maintain acetyl-CoA flux beyond a few hours. The contribution of protein deacetylation to acetyl-CoA metabolism, however, has not been directly tested.

**Fig 1.**
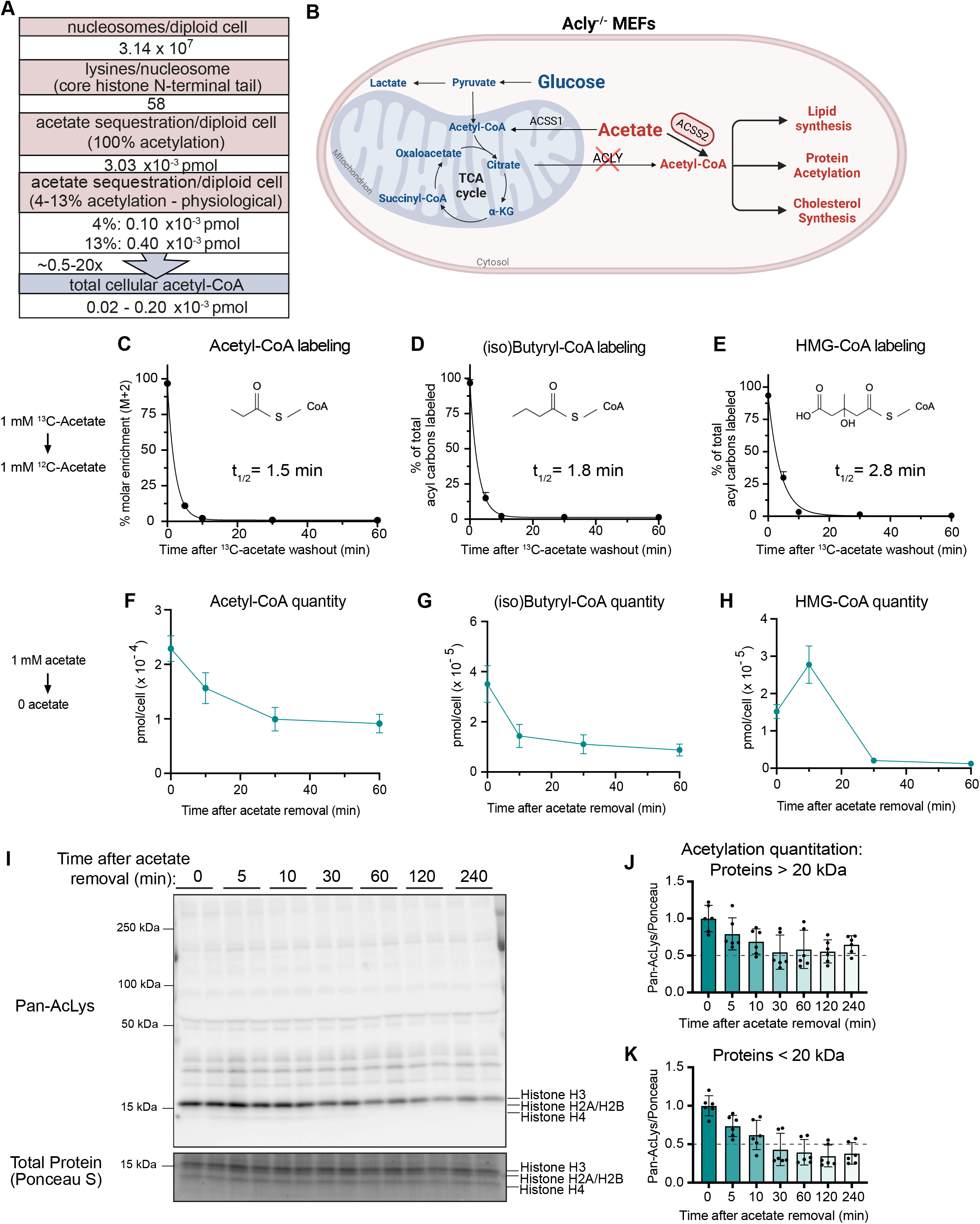
Acyl-CoA pools and protein acetylation respond rapidly to acetate availability in *Acly*^*-/-*^ MEFs. **(A)** Estimates of total and physiological acetate sequestration per human diploid cell compared to total acetyl-CoA pool size. Based on calculations in S1 Table. **(B)** Schematic representation of compartmentalized glucose and acetate metabolism in *Acly*^*-/-*^ MEFs. **(B-D)** Half-life measurement of **(C)** acetyl-CoA, **(D)** (iso)butyryl-CoA, **(E)** HMG-CoA in *Acly*^*-/-*^ MEFs cultured in 1mM acetate. N=9 across three independent experiments. **(F-H)** Whole-cell pool size measurement of **(F)** acetyl-CoA, **(G)** (iso)butyryl-CoA, **(H)** HMG-CoA at indicated time points after acetate removal. N=9 across three independent experiments. **(I)** Representative immunoblot of pan-acetyl-lysine in whole cell extracts upon 1mM acetate removal. **(J)** Quantification of pan-acetyl lysine more than 20 kilodaltons (kDa) upon 1mM acetate removal, normalized to total protein stain signal more than 20 kDa (Ponceau S). N=6 across three independent experiments.**(K)** Quantification of pan-acetyl lysine signal less than 20 kDa, which represents histone acetylation (H3 = 15 kDa; H2A/H2B = 14 kDa; H4 = 11 kDa). N=6 across three independent experiments. All data indicated as mean +/-standard deviation. Panel B adapted from “Resting metabolic activity vs stimulated metabolic activity” by BioRender.com (2022). Retrieved from https://app.biorender.com/biorender-templates

**Fig 2.**
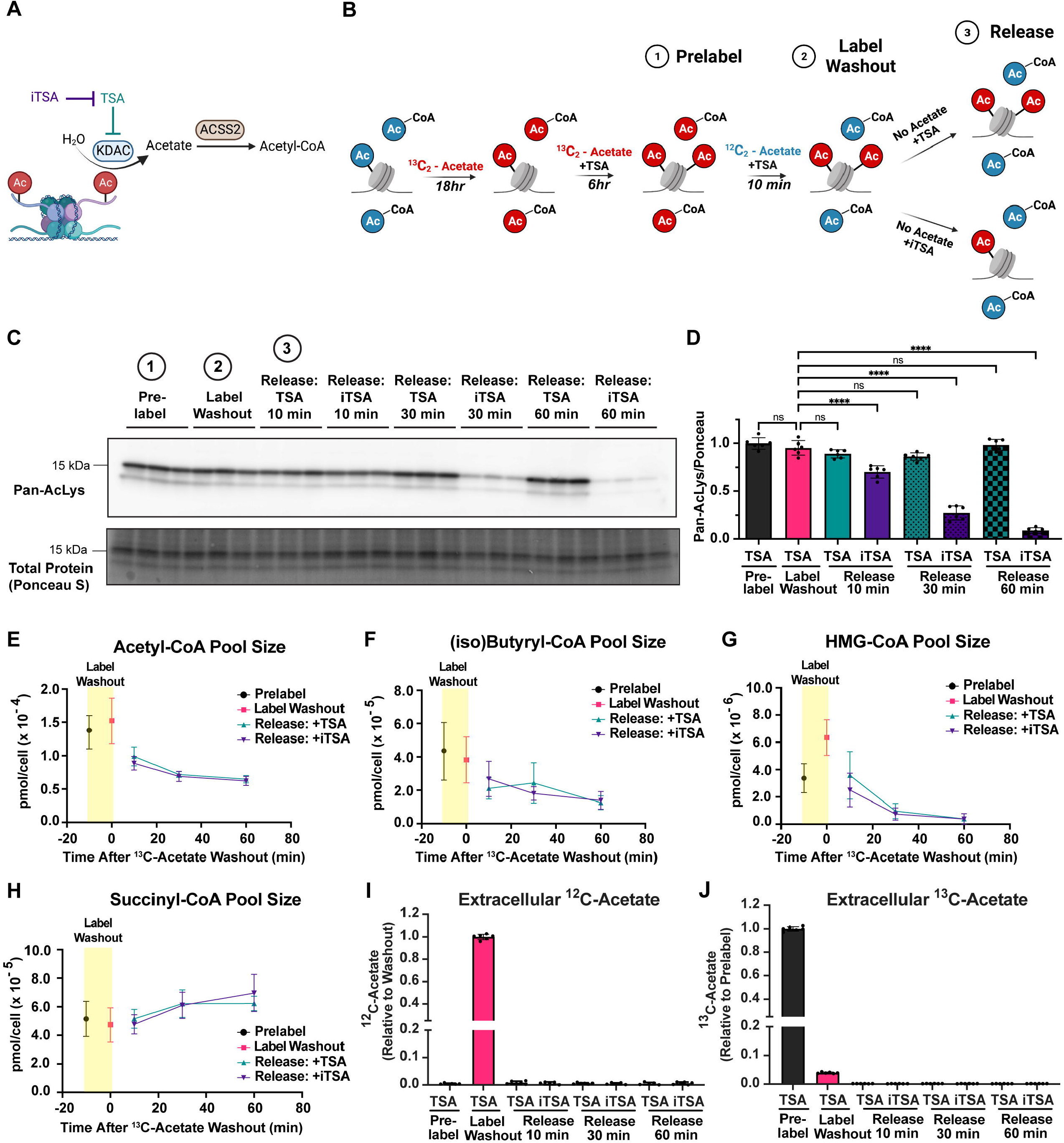
Protein deacetylation does not substantially impact cellular acyl-CoA levels. **(A)** Schematic representation of histone deacetylation contributing to acetyl-CoA. **(B)** Diagram illustrating protein acetylation prelabel, washout and release steps. Only histone acetylation is shown for simplicity. **(C)** Representative pan-acetyl-lysine immunoblot of whole cell extracts during prelabel, washout and release steps. **(D)** Quantification of pan-acetyl lysine signal below 20 kDa, normalized to total protein stain (Ponceau S) below 20 kDa. N=6 across two independent experiments. **(E)** Amount of whole-cell acetyl-CoA during prelabel, washout and release. **(F-H)** Same as in **(E)** but for **(F)** (iso)butyryl-CoA, **(G)** HMG-CoA and **(H)** succinyl-CoA. N=7 across two independent experiments. **(I-J)** Measurement of extracellular **(I)** ^12^C-acetate and **(J)** ^13^C-acetate under prelabel, washout and release conditions. N=6 across two independent experiments. All data indicated as mean +/-standard deviation. In **(D)**, a one-way ANOVA followed by Dunnett’s Multiple Comparison test was used to determine statistical significance. ns = not significant. P-value <0.05 (*), p-value < 0.01 (**), p-value <0.001 (***), P-value < 0.0001 (****).

### Acyl-CoA pools and protein acetylation respond rapidly to acetate availability in *Acly*^*-/-*^ MEFs

Manipulating and tracing histone acetylation marks is challenging because acetyl-CoA can be generated from a wide variety of metabolic sources. To overcome this problem, we used a well-characterized cell line model in which the metabolic source of acetyl-CoA was restricted. Mouse embryonic fibroblasts deficient in ATP citrate lyase (*Acly*^*-/-*^ MEFs) are unable to efficiently use mitochondrially-derived citrate to synthesize acetyl-CoA and are thus primarily dependent on acetate for cytosolic and nuclear acetyl-CoA, which supports fatty acid synthesis and histone acetylation [19] (Fig 1B). Although glucose is the primary source of mitochondrial citrate and tricarboxylic acid cycle intermediates in these cells, mitochondrial acetyl-CoA does not make a significant quantitative contribution to whole cell levels of acetyl-CoA [30,41]. Critically, bulk histone acetylation in *Acly*^*-/-*^ MEFs is sensitive to acetate supply: total histone lysine acetylation was lower in *Acly*^*-/-*^ MEFs compared to wildtype counterparts at physiological acetate concentration (100μM), but upon incubation with supraphysiological acetate (1mM), histone acetylation levels were increased back to wildtype levels [19]. Furthermore, the vast majority of histone acetylation (>80%) in *Acly*^*-/-*^ MEFs comes from acetate at supraphysiological acetate (see S1 Table for specific lysine sites) [19]. We therefore reasoned that *Acly*^*-/-*^ MEFs at supraphysiological acetate afforded us the unique opportunity both to manipulate the level of histone acetylation and to efficiently label acetylation marks with just one substrate.

We first tested the dynamics of acetate flux though acetyl-CoA and downstream products. *Acly*^*-/-*^ MEFs were incubated in 1mM ^13^C_2_-acetate for 16 hours to achieve maximal steady state labeling of acetyl-CoA >90%, then switched to media containing 1mM unlabeled acetate (Fig 1C). The rate of label loss for acetyl-CoA occurred with a half-time of 1.5 minutes. We also observed this turnover propagated downstream into abundant nucleocytosolic acyl-CoAs (iso)butyryl-CoA (4 carbon acyl chain) and HMG-CoA (6-carbon acyl chain) with half-times of 1.8 minutes and 2.8 minutes, respectively (Fig 1C-E). These turnover rates show that acetate flux through acetyl-CoA and downstream pathways is rapid, and that acetyl-CoA is a pacemaker for these downstream pathways, in agreement with previous kinetic studies [40].

Since these metabolites are highly dependent on acetate, we next asked how rapidly the pool sizes of these metabolites would respond to acetate removal. Consistent with their rapid turnover, acetyl-CoA, (iso)Butyryl-CoA and HMG-CoA were depleted by more than 50% after only 30 minutes (Fig 1E-G). Intriguingly, label loss after incubation in 1mM ^13^C_2_-acetate slowed upon acetate removal, suggesting that acetate removal also slows the rapid turnover of acetyl-CoA and downstream metabolites (S1A-C Fig). Acetate removal also caused rapid protein deacetylation: phistone acetylation was depleted within only 30 minutes, mirroring acetyl-CoA pool sizes (Fig 1I-K). Together, these data indicate that *Acly*^*-/-*^ MEFs provided a uniquely pliable system for manipulating and tracing acetyl-CoA and histone acetylation marks.

### Protein deacetylation does not substantially impact cellular acyl-CoA quantity

To determine the potential contribution of histone acetylation to acyl-CoA pools, we developed a modified pulse-chase experimental system in *Acly*^*-/-*^ MEFs in which we first maximize the amount of labeled histone acetylation with ^13^C_2_-acetate, then wash out ^13^C label from free acyl-CoAs while maintaining the ^13^C label on protein acetylation by inhibiting deacetylation, and finally release labeled ^13^C from protein in the absence of exogenous acetate (Fig 2A-B). In this experimental system, any appearance of ^13^C label in acyl-CoA pools after release of deacetylation likely derives from protein-bound acetate.

We first established the optimal conditions for each step in this experiment. We incubated *Acly*^*-/-*^ MEFs in 1 mM ^13^C_2_-acetate, which was previously shown to achieve on average greater than 80% labeling of histone acetylation marks [19]. To maximize labeling of protein acetylation, we sought to inhibit deacetylation. Class I and II KDACs appeared to be the main regulators of bulk deacetylation in this system since inhibitors (nicotinamide and sirtinol) of sirtuins (Class III KDAC) had no major impact on histone acetylation when used at concentrations previously shown to inhibit sirtuin activity [42–44] (S2A-B Fig). Sirtuin-mediated deacetylation produces O-Acetyl-ADP-ribose, which needs to be further metabolized to acetate (rather than directly generating acetate [45,46]), further suggesting sirtuin activity is not a major contributor to protein-derived acetate. Multiple Class I and II KDAC inhibitors, however, rapidly increased histone acetylation (S2B Fig). We found that the well-characterized Class I and II KDAC inhibitor, trichostatin A (TSA) [47,48] led to maximal acetylation after 6 hours in *Acly*^*-/-*^ MEFs (S3A Fig) and thus decided to use it to inhibit deacetylation in our system.

To wash the ^13^C label out of acyl-CoA intermediates we switched from 1mM ^13^C_2_-acetate to 1mM unlabeled acetate in the presence of TSA. ^13^C label was effectively washed out from abundant short chain acyl-CoA species within 10 min (Fig 3B-E), consistent with their fast turnover (Fig 1C-E).

**Fig 3.**
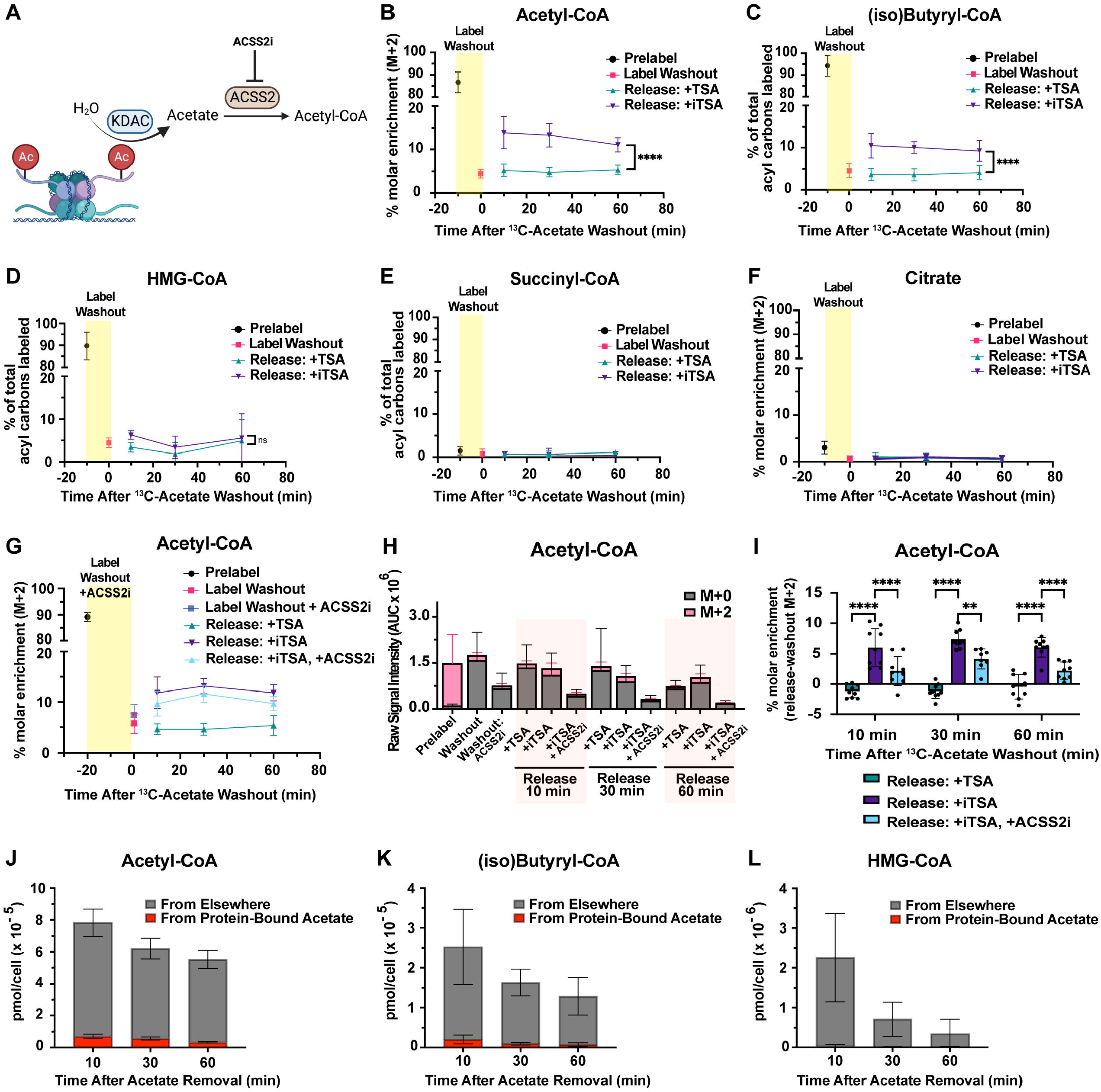
Protein deacetylation contributes carbons to acetyl-CoA and downstream pathways. **(A)** Schematic representation of histone deacetylation contributing to acetyl-CoA. **(B)** Percent molar enrichment of acetyl-CoA during prelabel, washout and release steps. **(C)** Percent of acyl carbons labeled of (iso)butyryl-CoA upon prelabel, washout and release. **(D-E)** Same as in (**B)** but for **(C)** HMG-CoA and **(D)** succinyl-CoA. N>9 across at least three independent experiments. **(F)** Same as in **(A)** but for citrate. N=6-7 across two independent experiments. **(G)** Percent molar enrichment of acetyl-CoA after prelabel, washout and release steps with ACSS2i (22μM) added as indicated. N=9 across three independent experiments. **(H)** Raw and uncorrected area under the curve (AUC) of acetyl-CoA unlabeled (M+0) and M+2 isotopologues after prelabel, washout and release with ACSS2i (22μM) added as indicated. N=9 across three independent experiments. **(I)** Percent molar enrichment from **(G)** of acetyl-CoA at 10, 30 and 60 minutes of release with washout enrichment subtracted. N=9 across three independent experiments. **(J)** Amount (pmol/cell) of acetyl-CoA that is derived from protein-bound acetate or elsewhere at 10, 30 or 60 minutes after acetate removal in ACLY^-/-^ MEFs. Calculated by multiplying percent molar enrichment by pool size at each timepoint. **(K-L)** Same as in **(J)** but for **(K)** (iso)butyryl-CoA and **(L)** HMG-CoA. All data indicated as mean +/-standard deviation. For **(B)-(D)** a two-way ANOVA followed by Sidak’s multiple comparison test was used to determine statistical significance. For **(I)** a two-way AVOVA followed by Dunnett’s Multiple Comparison was used to determine statistical significance.ns = not significant. P-value <0.05 (*), p-value < 0.01 (**), p-value <0.001 (***), P-value < 0.0001 (****).

To induce deacetylation, we then removed TSA and acetate from the cell culture medium. We at first observed that the deacetylation rate was much slower following TSA treatment (S3B Fig). We reasoned this may reflect persistent TSA activity. To promote rapid deacetylation after TSA treatment, we tested the efficacy of an inhibitor of TSA (iTSA) [49]. Treatment with iTSA led to deacetylation in minutes instead of hours (S3C Fig). We observed no significant difference in protein acetylation between prelabel, washout and release in the presence of TSA, consistent with TSA effectively preventing deacetylation. iTSA significantly decreased protein acetylation within 10 minutes of acetate removal (Fig 2C-D), similar to deacetylation dynamics observed with no TSA treatment (Fig 1K).

These experiments led us to establish the washout experimental system that consisted of three steps: 1) **Prelabel:** after 18 hour incubation with 1mM ^13^C_2_-acetate followed by 6 hours of TSA treatment were used to maximally load protein with labeled acetate; 2) **Washout:** 10 minutes of TSA and 1mM unlabeled acetate to wash out the label from acyl-CoAs but maintain it on protein-bound acetate; 3) **Release:** acetate removal and iTSA treatment to promote deacetylation (compared to a no release control with acetate removal in the presence of TSA) (Fig 2B).

Surprisingly, despite the significant capacity for acetate sequestration on protein, we found no significant difference in total quantity of cellular acetyl-CoA or other abundant short chain acyl-CoA species between deacetylation and controls (Fig 2E-H). Previous reports have shown that intracellular acetate can be acutely released from inside the cell into the extracellular space [13,21]. However, we found no detectable unlabeled acetate or ^13^C_2_-acetate in the cell culture medium after 10, 30 or 60 minutes of release (Fig 2I-J). A small amount of extracellular ^13^C_2_-acetate was detected in the label washout step likely because cells were not washed prior to switching to unlabeled washout media. Thus, we concluded that release of protein-derived acetate into the extracellular space was not a major fate of protein-bound acetate groups after induction of deacetylation.

### Protein deacetylation contributes carbons to acetyl-CoA and downstream pathways

We next investigated the potential for release of protein-bound acetate to contribute to flux of acyl-CoA pools. To do this, we compared the enrichment of ^13^C labeling in short chain acyl-CoAs in the prelabel, washout and release steps. As expected, approximately 90% of acyl-CoA pools were labeled at the prelabel step, and <5% of label remained after label washout (shaded yellow boxes, Figure 3B-D). The remaining label after washout is consistent with residual labeled acetate in the media during the washout step (Fig 2J).

The release step revealed a significant difference in the contribution of ^13^C labeling to acetyl-CoA between deacetylation and no release controls. During deacetylation (+iTSA), 15% of cellular acetyl-CoA was labeled after 10 minutes of acetate removal compared to less than 5% in no release (+TSA) controls. Enhanced labeling of acetyl-CoA persisted for at least 1-hour post-release, slowly declining to 10% enrichment (Fig 3B). We observed a similar response for (iso)butyryl-CoA labeling (Fig 3C). HMG-CoA labeling was marginally increased with deacetylation only at 10 minutes (Fig 3D). Consistent with the previous observation that acetate was not a major contributor to mitochondrial acetyl-CoA in *Acly*^*-/-*^ MEFs, there was less than 5% labeling of citrate and succinyl-CoA after the prelabel step and no difference upon induction of deacetylation (Fig 3E-F). Together, these data demonstrate that protein-derived acetate contributes carbons to acetyl-CoA and proximal downstream metabolites when deacetylation is induced by acetate removal.

The conversion of acetate to acetyl-CoA in the nuclear and cytosolic compartments is largely driven by ACSS2 [19,21]. To test if appearance of label in the acetyl-CoA pool was dependent on ACSS2 enzymatic activity, ACSS2 inhibitor (ACSS2i) was added during the label washout step (but not during the prelabel step because ACSS2 activity is required for prelabel). We extended the label washout from 10 to 20 minutes to allow sufficient time for inhibition. The washout step was still effective with ACSS2i, although there was a slight reduction in label washout efficiency (Fig 3G). The amount (i.e. raw signal detected) of acetyl-CoA was dramatically decreased with ACSS2i, indicating effective inhibition (Fig 3H). ACSS2i significantly reduced the appearance of label in acetyl-CoA between washout and release steps (Fig 3I), and a similar pattern was observed for (iso)butyryl-Coa (S4A-C Fig). These data demonstrate that ACSS2 enzymatic activity is required for protein-derived acetate incorporation into acetyl-CoA and downstream metabolites.

Finally, we calculated the quantity of protein-derived acetate that was incorporated into acetyl-CoA by combining isotope labeling experiments and pool size measurements at each timepoint. Deacetylation contributed 7.1x10^−6^ pmol/cell of acetyl-CoA 10 minutes after acetate removal, which made up approximately 9% of the total acetyl-CoA pool (Fig 3J). By 60 minutes of acetate removal, 3.4x10^−6^ pmol/cell of acetyl-CoA was derived from protein-bound acetate, approximately 6% of the acetyl-CoA pool, since both the pool size and labeling were reduced over the timecourse. A similar pattern was observed for (iso)butyryl-CoA: 2x10^−6^ pmol/cell of (iso)butyryl-CoA were derived from protein-bound acetate after 10 minutes of acetate removal, which decreased to 7.6x10^−7^ pmol/cell after 60 minutes (Fig 3K). Finally, only about 1% of the HMG-CoA pool was derived from protein-bound acetate after 10 minutes of acetate removal, likely due to inhibition of HMG-CoA synthesis upon acetate withdrawal (Fig 3L). Although we cannot confidently calculate total flux over the 60-minute timecourse in this non-steady state system, the maximum quantity of protein-derived acetyl-CoA detected at 10 minutes amounts to ∼0.3% of the maximal acetate we calculated that core histones could hold (Fig 1A).

## Discussion

We directly tested and quantified the potential for protein deacetylation to contribute to cellular acetyl-CoA by leveraging *Acly*^*-/-*^ MEFs, which provide a uniquely pliable system for manipulating and tracing acetyl-CoA and lysine acetylation. After inducing deacetylation of maximally acetylated protein, we detect 2-carbon labeling from protein-derived acetate in approximately 9% of cellular acetyl-CoA. Consistent with direct enzymatic conversion of free acetate to acetyl-CoA, 2-carbon labeling into acetyl-CoA was sensitive to ACSS2 inhibition. We also observed label incorporation into 4-carbon units of (iso)butyryl-CoA, suggesting that deacetylation-derived carbons can be propagated into downstream metabolites.

### Protein deacetylation is an acute carbon source for acetyl-CoA

Our data suggest that protein deacetylation is a short-lived contributor to acetyl-CoA metabolism, on the order of minutes. We detected the highest contribution of protein deacetylation to acetyl-CoA at the shortest timepoint (10 minutes after deacetylation induction) and this contribution decreased over the 60 minutes of deacetylation monitored. Our protein acetylation data show that deacetylation occurs predominantly between 10 and 30 minutes after acetate removal (Fig 2C, S2B Fig), indicating that deacetylation contribution was likely not more substantial prior to 10 minutes. Despite contributing to acetyl-CoA pools, deacetylation did not affect the total cellular concentration of acetyl-CoA upon acetate removal. Our data thus suggest that even though 50% of the acetyl-CoA pool is sustained for at least an hour in the absence of acetate and ACLY (Fig 1F), this is a result of decreased turnover (S1 Fig) and possible contribution from alternative metabolic source(s) other than deacetylation [50]. The functional consequence of deacetylation as an acute carbon source of acetyl-CoA is unclear. Future studies testing the effect on viability, differentiation or other functional outputs in cell types likely to be exposed to highly variable nutrient supply (such as endothelial or cancer cells) would be of interest.

### Protein deacetylation can potentially support acetyl-CoA-dependent anabolic pathways

We find that HMG-CoA levels drop by more than an order of magnitude 30 minutes after acetate removal in *Acly*^*-/-*^ MEFs, while succinyl-CoA levels are maintained (Fig 2G,H). Acetate removal therefore shuts down HMG-CoA-dependent anabolic pathways such as the mevalonate pathway in *Acly*^*-/-*^ MEFs but cells maintain TCA cycle-dependent catabolism (as reported previously [40]). Nevertheless, we find that protein-derived acetate contributes carbon units to (iso)butyryl-CoA in addition to acetyl-CoA. We identify (iso)butyryl-CoA as downstream of acetyl-CoA in our system, since labeling occurs rapidly upon ^13^C-acetate introduction but is slightly slower than acetyl-CoA (Fig 1C,D), consistent with pattern observed in other downstream pathways [40]. Furthermore, ACSS2 inhibition reduces carbon contribution of protein-derived acetate to (iso)butyryl-CoA (S4 Fig). Importantly, (iso)butyryl-CoA can be an intermediate in the fatty acid synthesis pathway downstream of acetyl-CoA [51]. Our data therefore suggest the intriguing possibility that protein-derived acetate can be used for anabolic pathways such as fatty acid synthesis. Future studies will illuminate the quantitative contribution of protein-derived acetate to fatty acid synthesis and other anabolic pathways under conditions and in cell types where anabolism is activated.

Histones are highly abundant proteins and likely account for a majority of the protein acetylation in the cell [34,37–39] (S2 Table), which suggests that protein deacetylation may specifically enrich the nuclear compartment with protein-derived acetate [21]. Therefore, even though TSA is a Class I and II KDAC inhibitor that inhibits both nuclear and cytosolic/mitochondrial KDACs, the availability of protein-derived acetate for non-nuclear uses may be limited because the acetylation stoichiometry of non-nuclear compartments is minimal and the nucleus is a distinct metabolic compartment [21,30,52,53]. Measurements of the relative contribution of protein deacetylation to acetyl-CoA and downstream processes within specific subcellular compartments would be required to investigate the potential for compartment-specific functions of protein/histone deacetylation.

### Role of dynamic histone acetylation in gene regulation

The dynamic response of histone acetylation to acetate availability in *Acly*^*-/-*^ MEFs emphasizes the dynamic nature of post-translational lysine acetylation, which has been previously shown to have a half-life on the order of minutes [18,20]. Histone lysine acetylation is an epigenetic mark that is believed to promote gene expression through at least two mechanisms: 1) promoting chromatin decompaction by neutralizing lysine positive charge on histones [54], and 2) through specific interactions recruiting transcriptional machinery [55]. Whether bulk acetylation/deacetylation in response to acetate supply in the *Acly*^*-/-*^ MEFs model preferentially affects specific histone lysine residues or specific genome regions is an interesting question for future investigation. Indeed, differential sensitivity of specific histone lysine sites to acetyl-CoA has been identified in mammalian cells [6,56]. Furthermore, genome-wide studies of histone acetylation dynamics in yeast revealed that different histone lysine residues exhibited distinct responses to quiescence exit even though global histone acetylation increased overall [23]. Intriguingly, certain histone lysine sites appear to function as acetate reservoirs, while others are site-specifically regulated to mediate rapid changes in transcription [23]. Future studies coupling measurements of compartment-specific contribution of protein deacetylation with chromatin-wide acetylation patterns would give added insight as to where the deacetylation-derived acetate is incorporated, and begin to shed light on why it has limited potential for maintaining cellular acetyl-CoA-dependent metabolic pathways.

In summary, we demonstrate here that protein deacetylation can be an acute carbon source to generate acetyl-CoA, transiently supplying approximately 9% of the acetyl-CoA pool after maximal acetylation. Surprisingly, this contribution represents <1% of potential contribution we and others estimate to be on maximally acetylated histones. Collectively, this work presents a framework to examine the interplay between the epigenetic and metabolic functions of protein acetylation.

## Materials and methods

### Cell culture

*Acly*^*-/-*^ MEFs were generated in the Wellen lab as reported previously [19]. *Acly*^*-/-*^ MEFs were maintained at <80% confluence (passaged every 3-4 days) in high glucose DMEM (Gibco #11965-084) with 10% fetal bovine serum (FBS, Hyclone cat# SH30071.03, lot# AF29485596). *Acly*^*-/-*^ MEFs were negative for mycoplasma, tested through the University of Pennsylvania Cell Center Mycoplasma Testing Service.

### Washout experimental system

For all washout experiments including stable isotope tracing, *Acly*^*-/-*^ MEFs in 6cm dishes at 60-80% confluence were washed with phosphate buffered saline (PBS, Corning #21-040-CM) and switched from DMEM + 10% FBS to DMEM + 10% dialyzed FBS (dFBS, GeminiBio #100-108) + 1mM ^13^C_2_ - acetate (Cambridge Isotope Laboratories #CLM-440-0) for 18 hours, so as to label protein acetylation [19]. Prelabel with trichostatin A (TSA) was done by aspirating medium and switching to DMEM + 10% dFBS + 1mM ^13^C_2_ – acetate + 300 nM TSA (Selleck Chemicals #S1054) for 6 hours (note no PBS wash). Label was removed from intracellular metabolites but maintained on protein by aspirating labeled TSA medium and replacing with unlabeled TSA medium, i.e. DMEM + 10% dFBS + 1mM ^12^C – acetate (Sigma #S7547) + 300 nM TSA for 10 min (no PBS wash). Finally, protein acetylation was released by washing with PBS and replacing medium with DMEM + 10% dFBS + 300nM TSA as control group or DMEM + 10% dFBS + 50 μM iTSA1 (Cayman Chemical #18681) as release group. At least three 6cm dishes per condition were collected by direct extraction (described below) after TSA preloading, after label washout, and after 10-60 min of release. Unlabeled control dishes were incubated overnight in DMEM + 10% dialyzed FBS + 1mM ^12^C – acetate, media was changed as described above but with ^12^C-acetate instead of ^13^C-acetate and collected during release step; these unlabeled dishes were used to correct the fractional enrichment for natural isotope abundance within each experiment using FluxFix [57].

When ACSS2 inhibitor (Selleck #S8588, CAS No. 508186-14-9) was added, the experiment above was altered in three ways: 1) the label washout period was increased to 20 minutes; 2) an additional label washout group was included that consisted of DMEM + 10% dFBS + 1mM ^12^C – acetate + 300 nM TSA + 22μM ACSS2i; and 3) an additional release group was included consisting of DMEM + 10% dFBS + 50 μM iTSA1 + 22μM ACSS2i.

For acyl-CoA pool size quantification in the washout experimental paradigm depicted in Figure 2, the same experimental strategy described above was employed, except that ^12^C-acetate was used for all steps requiring acetate. In addition, at least one dish per experimental condition was used for quantifying cell number (Corning #6749) and packed cell volume (TPP #87005).

### Whole cell extraction for metabolite processing

Direct extraction of whole cell metabolites was carried out as described previously[40]. For short-chain acyl-CoA detection, media was poured into a waste container, residual media was subsequently aspirated and dishes were placed on ice. 1ml of ice-cold 10% (w/v in water) TCA (Sigma #T6399) was quickly added to each dish. Cells were scraped, transferred into 1.5 ml tubes on ice, and frozen at -80°C until further processing.

For citrate detection, 80:20 methanol:water extraction was used. Media was quickly aspirated, cells were washed with 2ml ice-cold PBS and then 1ml of 80:20 HPLC grade methanol:water (Optima) cooled at -80°C was added. Cells were scraped while on dry ice, transferred into 1.5 ml tubes and frozen at -80°C until further processing.

For acyl-CoA pool size quantification, direct 10% TCA extraction was used to quench cell metabolism, and samples were then spiked with 0.1 mL of ^13^C_3_^15^N_1_ -acyl-CoA internal standard prepared from yeast as previously described[58]. Cells were then scraped, transferred into 1.5 ml tubes on ice, and frozen at -80°C until further processing. Calibration curves were prepared from commercially available acyl-CoA standards (Sigma Aldrich). Metabolite pool size was normalized to cell number.

### Immunoblotting

Immunoblots were conducted on protein precipitated after direct 10% TCA extraction. Samples were thawed from -80°C and protein pellets were spun down at 13,000 x g, 10 minutes at 4°C. Pellets were washed with 1mL of acetone, spun down again, then acetone was decanted and pellet was allowed to dry. Pellets were then resuspended in 80μl 2% SDS in 10mM Tric-HCL pH 7.5 and vortexed/incubated at 37°C until dissolved. Supernatant were taken after further centrifugation at 10,000 x g, 5 minutes at room temperature to eliminate insoluble matter. Protein content was measured with Pierce BCA Protein Assay Kit (Thermo Scientific #23225), adjusted to equal concentration for each sample and boiled at 95°C for 10 minutes in Laemmli buffer (Bio-Rad #161-0747).

Samples (20-30μg of protein) were run on a 4-20% gradient tris-polyacrylamide gel (Bio-Rad #3450034) at 100 volts for 1.25 hours. Protein was transferred onto PVDF membranes (Millipore #IPVH00010), which were then incubated with Ponceau (0.1% w/v Ponceau S (Sigma #P3504) in 5% acetic acid) for 10 minutes. Ponceau-stained membranes were imaged on a digital imager (GE ImageQuant LAS 4000) on digitization (epi-illumination) mode. Membranes were then blocked in 5% non-fat dry milk in tris-buffered saline, 0.1%-tween (TBST) for 30 minutes and incubated in primary antibody overnight. The next day, membranes were washed with TBST for 10 minutes 3 times, then incubated for 60 minutes in rabbit horseradish peroxidase-conjugated secondary antibody (1:10,000 Cell Signaling Technology #7074) and washed again with TBST 3 times, 10 minutes each. Enhanced chemiluminescent HRP substrate (Thermo #34095) was used to determine protein signal on a digital imager (GE LAS 4000). Pan-acetyl lysine primary antibody (ICP #0380) was used for Supp. Figure 1C, and pan-acetyl lysine primary antibody (CST #9441) was used for all other immunoblots.

Immunoblots were quantified with Fiji software [59]. Mean gray value for each pan-acetyl-lysine lane was determined, and then mean gray value for an empty lane was used as background subtraction. Each background subtracted value was then divided by that sample’s specific ponceau signal (also mean gray value) to account for loading variability.

### Sample processing for liquid-chromatography mass spectrometry

Samples were thawed and kept on ice throughout processing. Cell and fraction samples in 10% (w/v) trichloroacetic acid in water were sonicated for 12 × 0.5 s pulses, protein was pelleted by centrifugation at 13,000 ×*g* from 10 min at 4 °C. The supernatant was purified by solid-phase extraction using Oasis HLB 1cc (30 mg) SPE columns (Waters). Columns were washed with 1 mL methanol, equilibrated with 1 mL water, loaded with supernatant, desalted with 1 mL water, and eluted with 1 mL methanol containing 25 mM ammonium acetate. The purified extracts were evaporated to dryness under nitrogen then resuspended in 55 μl 5% (w/v) 5-sulfosalicylic acid (SSA) in water.

### Acyl-CoA analysis by liquid-chromatography mass spectrometry

Acyl-CoAs were measured by liquid chromatography-high resolution mass spectrometry. 5–10 μl of purified samples in 5% SSA were analyzed by injection of an Ultimate 3000 Quaternary UHPLC coupled to a Q Exactive Plus (Thermo Scientific) mass spectrometer in positive ESI mode using the settings described previously [60]. Quantification of acyl-CoAs was via their [M+H]+ ions, and isotope tracing via their MS2 product ion from the predominant [M+H-507]+ neutral loss fragment with the targeted masses used for isotopologue analysis are indicated in Table 1. Data were integrated using Tracefinder v4.1 (Thermo Scientific) software. Isotopic enrichment in tracing experiments was calculated by normalization to unlabeled control samples using the FluxFix calculator [57].

**Table 1:**
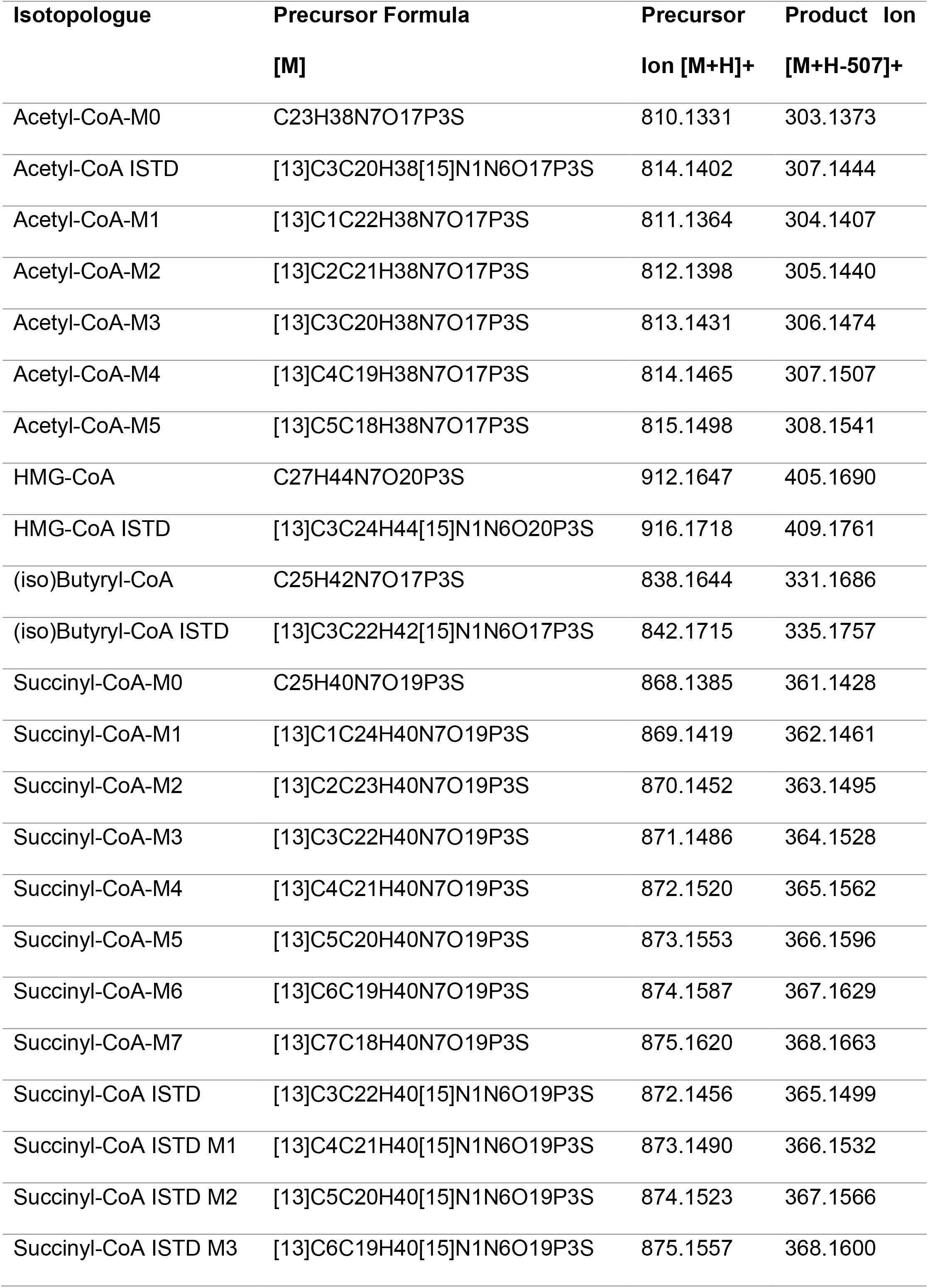
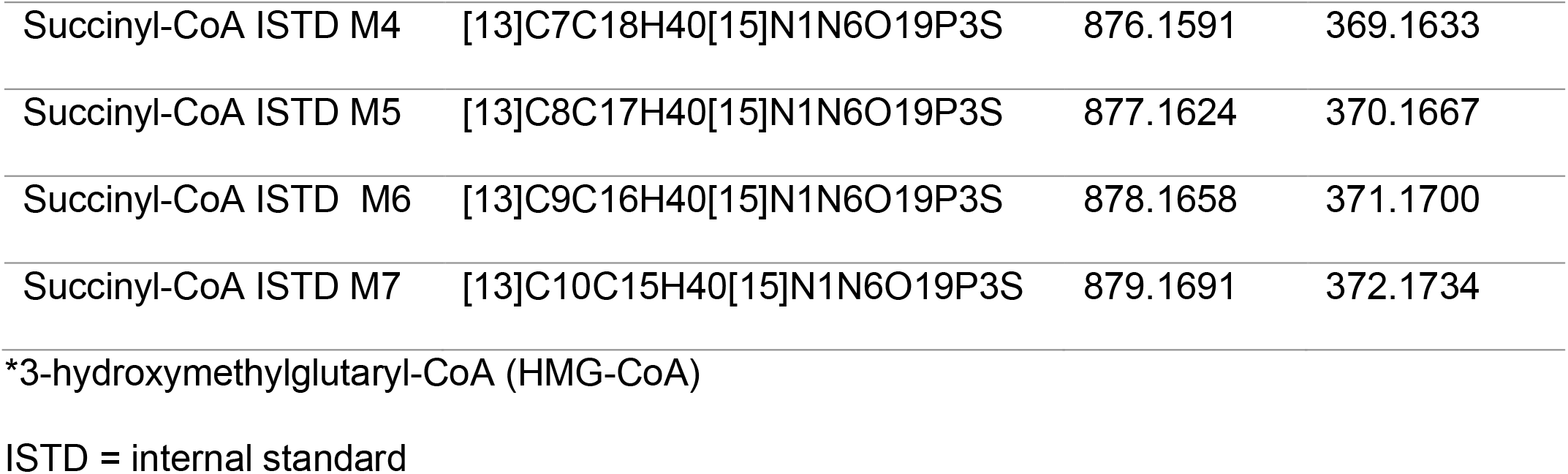
Acyl-CoA Masses.

| **Isotopologue** | **Precursor Formula**<br>**[M]** | **Precursor**<br>**Ion [M+H]<sup>+</sup>** | **Product Ion**<br>**[M+H-507]<sup>+</sup>** |
| --- | --- | --- | --- |
| Acetyl-CoA-M0 | C23H38N7O17P3S | 810.1331 | 303.1373 |
| Acetyl-CoA ISTD | [13]C3C20H38[15]N1N6O17P3S | 814.1402 | 307.1444 |
| Acetyl-CoA-M1 | [13]C1C22H38N7O17P3S | 811.1364 | 304.1407 |
| Acetyl-CoA-M2 | [13]C2C21H38N7O17P3S | 812.1398 | 305.1440 |
| Acetyl-CoA-M3 | [13]C3C20H38N7O17P3S | 813.1431 | 306.1474 |
| Acetyl-CoA-M4 | [13]C4C19H38N7O17P3S | 814.1465 | 307.1507 |
| Acetyl-CoA-M5 | [13]C5C18H38N7O17P3S | 815.1498 | 308.1541 |
| HMG-CoA | C27H44N7O20P3S | 912.1647 | 405.1690 |
| HMG-CoA ISTD | [13]C3C24H44[15]N1N6O20P3S | 916.1718 | 409.1761 |
| (iso)Butyryl-CoA | C25H42N7O17P3S | 838.1644 | 331.1686 |
| (iso)Butyryl-CoA ISTD | [13]C3C22H42[15]N1N6O17P3S | 842.1715 | 335.1757 |
| Succinyl-CoA-M0 | C25H40N7O19P3S | 868.1385 | 361.1428 |
| Succinyl-CoA-M1 | [13]C1C24H40N7O19P3S | 869.1419 | 362.1461 |
| Succinyl-CoA-M2 | [13]C2C23H40N7O19P3S | 870.1452 | 363.1495 |
| Succinyl-CoA-M3 | [13]C3C22H40N7O19P3S | 871.1486 | 364.1528 |
| Succinyl-CoA-M4 | [13]C4C21H40N7O19P3S | 872.1520 | 365.1562 |
| Succinyl-CoA-M5 | [13]C5C20H40N7O19P3S | 873.1553 | 366.1596 |
| Succinyl-CoA-M6 | [13]C6C19H40N7O19P3S | 874.1587 | 367.1629 |
| Succinyl-CoA-M7 | [13]C7C18H40N7O19P3S | 875.1620 | 368.1663 |
| Succinyl-CoA ISTD | [13]C3C22H40[15]N1N6O19P3S | 872.1456 | 365.1499 |
| Succinyl-CoA ISTD M1 | [13]C4C21H40[15]N1N6O19P3S | 873.1490 | 366.1532 |
| Succinyl-CoA ISTD M2 | [13]C5C20H40[15]N1N6O19P3S | 874.1523 | 367.1566 |
| Succinyl-CoA ISTD M3 | [13]C6C19H40[15]N1N6O19P3S | 875.1557 | 368.1600 |
| Succinyl-CoA ISTD M4 | [13]C7C18H40[15]N1N6O19P3S | 876.1591 | 369.1633 |
| Succinyl-CoA ISTD M5 | [13]C8C17H40[15]N1N6O19P3S | 877.1624 | 370.1667 |
| Succinyl-CoA ISTD M6 | [13]C9C16H40[15]N1N6O19P3S | 878.1658 | 371.1700 |
| Succinyl-CoA ISTD M7 | [13]C10C15H40[15]N1N6O19P3S | 879.1691 | 372.1734 |

\*3-hydroxymethylglutaryl-CoA (HMG-CoA)
ISTD = internal standard

### Acetate quantification by gas-chromatography mass spectrometry

To harvest media for extracellular acetate measurement, 1mL of media was collected, spun down at 2500 x g for 5 minutes and frozen at -80°C until further processing. Acetate measurement was done by adapting a previously described gas-chromatography mass spectrometry method [37]. 200μL of media was added to a 2mL eppendorf tube followed by 40μL of 1mM ^13^C_2_-D_3_ sodium acetate (Cambridge Isotope #CDLM-3457) as internal standard. 50μL of 1-propanol and 50μL of pyridine were added, and samples were incubated on ice for 5 minutes. 100μL of 1M sodium hydroxide was then added, immediately followed by 30μL methyl chloroformate. Samples were vigorously vortexed for 20 seconds, pausing to vent halfway through. Finally, 300μL of methyl tert-butyl ether was added, samples were vortexed for 20 seconds without venting and centrifuged at 10,000 x g for 5 minutes at 4°C. 100-200μL of the upper layer was then transferred to gas-chromatography vials for analysis.

An Agilent 7890B gas chromatograph (GC) coupled with an Agilent 5977B mass selective detector (MSD) was used for analysis. The GC column was a 30 m x 250 μm x 0.25 μm HP-5ms Ultra Inert column (Agilent #19091S-433UI). 2μL of sample was injected at 15mL/min split flow (15:1 split ratio), with inlet temperature at 280°C. The oven was held at 45°C for 0.8 minutes, then ramped from 45°C to 60°C at 25°C/min and held for 0 minutes, then ramped from 60°C to 190°C at 50°C /min and held for 0 min. The mass spectrometer was operated in SIM mode for m/z 61 (^12^C-acetate), 63 (^13^C-acetate) and 66 (^13^C_2_; D_3_-acetate) with a dwell time of 50ms for each ion. Agilent MassHunter Qualitative Analysis software (B.07.00) was used for visualization of chromatograms and Agilent MassHunter Quantitative Analysis software was used for peak area integration. Peak area of m/z 61 and 63 for each sample was divided by m/z 66 to normalize to the internal standard. In each experiment, background ^12^C-acetate was subtracted out of all samples by using media blank, i.e. DMEM + 10% dFBS.

### Statistical analyses

All data are reported as mean ± standard deviation. To determine statistical significance between two or more groups, a one-way ANOVA followed by Dunnett’s Multiple Comparison Test was conducted. For comparing the mean of two groups over a time course (such as in Figure 3), a two-way ANOVA followed by Sidak’s multiple comparison test was used. Prism 9 (GraphPad) was used for graphing and statistical analyses, and Illustrator (Adobe) was used for figure layouts. Figure 1B, Figure 2A-B and Figure 3A were created with BioRender.com.

## Acknowledgments

We gratefully acknowledge conceptual and experimental input and discussion from Nora Yucel, and thank Arany and Wellen Lab members for feedback throughout the course of this project.

## Supporting Information

**S1 Fig. Acetate removal slows acyl-CoA turnover in *Acly***^***-/-***^ **MEFs. (A)** Percent molar enrichment (M+2) of acetyl-CoA after 1mM acetate removal. **(B)** Percent of total acyl carbons labeled of (iso)butyryl-CoA after 1mM acetate removal. **(C)** Same as in **(B)** but for HMG-CoA. N>6 across at least two independent experiments.

**S2 Fig. Class III KDAC inhibition has minimal effect on global histone acetylation. (A)** Immunoblot of pan-acetyl-lysine in acid extracted histones after 1mM acetate removal and Sirtinol (50μM), Apicidin (1μM), TSA (300nM), Panobinostat (400nM) or sodium butyrate (2mM) treatment at indicated timepoints. **(B)** Immunoblot of pan-acetyl-lysine in acid extracted histones after 1mM acetate removal and nicotinamide (5mM), sodium butyrate (2mM), Panobinostat (400nM), Apicidin (1μM) or TSA (300nM) treatment at indicated timepoints. 0.1mM and 1mM acetate represent no media change controls (beginning of timecourse).

**S3 Fig. iTSA speeds up reversal of TSA. (A)** Immunoblot of pan-acetyl-lysine after indicated timepoint of treatment with KDAC inhibitor TSA (300nM). **(B)** Immunoblot of pan-acetyl-lysine upon TSA removal after 10 minute TSA treatment (300nM) in the presence of 1mM acetate. **(C)** Immunoblot of pan-acetyl-lysine upon TSA removal with or without iTSA (50μM) after 6 hour TSA treatment (300nM) in the presence of 1mM acetate.

**S4 Fig. ACSS2i decreases (iso)butyryl-CoA pool size and isotopic labeling from acetate. (A)** Percent of total acyl carbons labeled of (iso)butyryl-CoA after prelabel, washout and release steps with ACSS2i (22μM) added as indicated. N=9 across three independent experiments. **(B)** Raw and uncorrected area under the curve (AUC) of (iso)butyryl-CoA unlabeled (M+0) and M+2 and M+4 isotopologues after prelabel, washout and release with ACSS2i (22μM) added as indicated. N=9 across three independent experiments. **(C)** % of total acyl carbons labeled from **(A)** of (iso)butyryl-CoA at 10, 30 and 60 minutes of release with washout enrichment subtracted. N=9 across three independent experiments. A two-way AVOVA followed by Dunnett’s Multiple Comparison was used to determine statistical significance. ns = not significant. P-value <0.05 (*), p-value < 0.01 (**), p-value <0.001 (***), P-value < 0.0001 (****). All data indicated as mean +/-standard deviation.

**S1 Table. Maximal and physiological acetate amount on N-terminal histone tails and acetylation occupancy of *Acly***^***-/-***^ **MEFs**.

**S2 Table. Distribution of acetylation across subcellular compartments**.

## References

1. Pietrocola F, Galluzzi L, Bravo-San Pedro JM, Madeo F, Kroemer G. Acetyl Coenzyme A: A Central Metabolite and Second Messenger. Cell Metabolism. 2015;21: 805–821. doi:10.1016/j.cmet.2015.05.014

2. Shi L, Tu BP. Acetyl-CoA and the regulation of metabolism: mechanisms and consequences. Curr Opin Cell Biol. 2015;33: 125–131. doi:10.1016/j.ceb.2015.02.003

3. Wellen KE, Hatzivassiliou G, Sachdeva UM, Bui TV, Cross JR, Thompson CB. ATP-citrate lyase links cellular metabolism to histone acetylation. Science. 2009;324: 1076–1080. doi:10.1126/science.1164097

4. Lee JV, Carrer A, Shah S, Snyder NW, Wei S, Venneti S, et al. Akt-dependent metabolic reprogramming regulates tumor cell histone acetylation. Cell Metab. 2014;20: 306–319. doi:10.1016/j.cmet.2014.06.004

5. Moussaieff A, Rouleau M, Kitsberg D, Cohen M, Levy G, Barasch D, et al. Glycolysis-Mediated Changes in Acetyl-CoA and Histone Acetylation Control the Early Differentiation of Embryonic Stem Cells. Cell Metabolism. 2015;21: 392–402. doi:10.1016/j.cmet.2015.02.002

6. Carrer A, Parris JLD, Trefely S, Henry RA, Montgomery DC, Torres A, et al. Impact of a High-fat Diet on Tissue Acyl-CoA and Histone Acetylation Levels. J Biol Chem. 2017;292: 3312–3322. doi:10.1074/jbc.M116.750620

7. Covarrubias AJ, Aksoylar HI, Yu J, Snyder NW, Worth AJ, Iyer SS, et al. Akt-mTORC1 signaling regulates Acly to integrate metabolic input to control of macrophage activation. eLife. 2016;5: e11612. doi:10.7554/eLife.11612

8. Finley LWS, Vardhana SA, Carey BW, Alonso-Curbelo D, Koche R, Chen Y, et al. Pluripotency transcription factors and Tet1/2 maintain Brd4-independent stem cell identity. Nat Cell Biol. 2018;20: 565–574. doi:10.1038/s41556-018-0086-3

9. Fernandez S, Viola JM, Torres A, Wallace M, Trefely S, Zhao S, et al. Adipocyte ACLY Facilitates Dietary Carbohydrate Handling to Maintain Metabolic Homeostasis in Females. Cell Reports. 2019;27: 2772-2784.e6. doi:10.1016/j.celrep.2019.04.112

10. Martinez Calejman C, Trefely S, Entwisle SW, Luciano A, Jung SM, Hsiao W, et al. mTORC2-AKT signaling to ATP-citrate lyase drives brown adipogenesis and de novo lipogenesis. Nat Commun. 2020;11: 575. doi:10.1038/s41467-020-14430-w

11. Kurdistani SK. Chromatin: a capacitor of acetate for integrated regulation of gene expression and cell physiology. Current Opinion in Genetics & Development. 2014;26: 53–58. doi:10.1016/j.gde.2014.06.002

12. Ye C, Tu BP. Sink into the Epigenome: Histones as Repositories That Influence Cellular Metabolism. Trends in Endocrinology & Metabolism. 2018;29: 626–637. doi:10.1016/j.tem.2018.06.002

13. McBrian MA, Behbahan IS, Ferrari R, Su T, Huang T-W, Li K, et al. Histone acetylation regulates intracellular pH. Mol Cell. 2013;49: 310–321. doi:10.1016/j.molcel.2012.10.025

14. Boon R, Silveira GG, Mostoslavsky R. Nuclear metabolism and the regulation of the epigenome. Nat Metab. 2020;2: 1190–1203. doi:10.1038/s42255-020-00285-4

15. Nirello VD, Rodrigues de Paula D, Araújo NVP, Varga-Weisz PD. Does chromatin function as a metabolite reservoir? Trends in Biochemical Sciences. 2022;47: 732–735. doi:10.1016/j.tibs.2022.03.016

16. Srivatsan SR, McFaline-Figueroa JL, Ramani V, Saunders L, Cao J, Packer J, et al. Massively multiplex chemical transcriptomics at single-cell resolution. Science. 2020;367: 45–51. doi:10.1126/science.aax6234

17. Simithy J, Sidoli S, Yuan Z-F, Coradin M, Bhanu NV, Marchione DM, et al. Characterization of histone acylations links chromatin modifications with metabolism. Nat Commun. 2017;8: 1141. doi:10.1038/s41467-017-01384-9

18. Evertts AG, Zee BM, Dimaggio PA, Gonzales-Cope M, Coller HA, Garcia BA. Quantitative dynamics of the link between cellular metabolism and histone acetylation. J Biol Chem. 2013;288: 12142–12151. doi:10.1074/jbc.M112.428318

19. Zhao S, Torres A, Henry RA, Trefely S, Wallace M, Lee JV, et al. ATP-Citrate Lyase Controls a Glucose-to-Acetate Metabolic Switch. Cell Reports. 2016;17: 1037–1052. doi:10.1016/j.celrep.2016.09.069

20. Weinert BT, Narita T, Satpathy S, Srinivasan B, Hansen BK, Schölz C, et al. Time-Resolved Analysis Reveals Rapid Dynamics and Broad Scope of the CBP/p300 Acetylome. Cell. 2018;174: 231-244.e12. doi:10.1016/j.cell.2018.04.033

21. Bulusu V, Tumanov S, Michalopoulou E, van den Broek NJ, MacKay G, Nixon C, et al. Acetate Recapturing by Nuclear Acetyl-CoA Synthetase 2 Prevents Loss of Histone Acetylation during Oxygen and Serum Limitation. Cell Reports. 2017;18: 647–658. doi:10.1016/j.celrep.2016.12.055

22. Li X, Yu W, Qian X, Xia Y, Zheng Y, Lee J-H, et al. Nucleus-Translocated ACSS2 Promotes Gene Transcription for Lysosomal Biogenesis and Autophagy. Molecular Cell. 2017;66: 684-697.e9. doi:10.1016/j.molcel.2017.04.026

23. Mendoza M, Egervari G, Sidoli S, Donahue G, Alexander DC, Sen P, et al. Enzymatic transfer of acetate on histones from lysine reservoir sites to lysine activating sites. Sci Adv. 2022;8: eabj5688. doi:10.1126/sciadv.abj5688

24. tan M, Luo H, Lee S, Jin F, Yang JS, Montellier E, et al. Identification of 67 Histone Marks and Histone Lysine Crotonylation as a New Type of Histone Modification. Cell. 2011;146: 1016–1028. doi:10.1016/j.cell.2011.08.008

25. Iwasaki W, Miya Y, Horikoshi N, Osakabe A, Taguchi H, Tachiwana H, et al. Contribution of histone N-terminal tails to the structure and stability of nucleosomes. FEBS Open Bio. 2013;3: 363–369. doi:10.1016/j.fob.2013.08.007

26. Routh A, Sandin S, Rhodes D. Nucleosome repeat length and linker histone stoichiometry determine chromatin fiber structure. Proc Natl Acad Sci U S A. 2008;105: 8872–8877. doi:10.1073/pnas.0802336105

27. Widom J. A relationship between the helical twist of DNA and the ordered positioning of nucleosomes in all eukaryotic cells. Proc Natl Acad Sci U S A. 1992;89: 1095–1099. doi:10.1073/pnas.89.3.1095

28. International Human Genome Sequencing Consortium. Finishing the euchromatic sequence of the human genome. Nature. 2004;431: 931–945. doi:10.1038/nature03001

29. Church DM, Goodstadt L, Hillier LW, Zody MC, Goldstein S, She X, et al. Lineage-specific biology revealed by a finished genome assembly of the mouse. PLoS Biol. 2009;7: e1000112. doi:10.1371/journal.pbio.1000112

30. trefely S, Huber K, Liu J, Noji M, Stransky S, Singh J, et al. Quantitative subcellular acyl-CoA analysis reveals distinct nuclear metabolism and isoleucine-dependent histone propionylation. Molecular Cell. 2022;82: 447-462.e6. doi:10.1016/j.molcel.2021.11.006

31. Abshiru N, Caron-Lizotte O, Rajan RE, Jamai A, Pomies C, Verreault A, et al. Discovery of protein acetylation patterns by deconvolution of peptide isomer mass spectra. Nat Commun. 2015;6: 8648. doi:10.1038/ncomms9648

32. Zhou T, Chung Y-H, Chen J, Chen Y. Site-Specific Identification of Lysine Acetylation Stoichiometries in Mammalian Cells. J Proteome Res. 2016;15: 1103–1113. doi:10.1021/acs.jproteome.5b01097

33. Zheng Y, Thomas PM, Kelleher NL. Measurement of acetylation turnover at distinct lysines in human histones identifies long-lived acetylation sites. Nat Commun. 2013;4: 2203. doi:10.1038/ncomms3203

34. Hansen BK, Gupta R, Baldus L, Lyon D, Narita T, Lammers M, et al. Analysis of human acetylation stoichiometry defines mechanistic constraints on protein regulation. Nat Commun. 2019;10: 1055. doi:10.1038/s41467-019-09024-0

35. Guo Q, Sidoli S, Garcia BA, Zhao X. Assessment of Quantification Precision of Histone Post-Translational Modifications by Using an Ion Trap and down To 50 000 Cells as Starting Material. J Proteome Res. 2018;17: 234–242. doi:10.1021/acs.jproteome.7b00544

36. Yucel N, Wang YX, Mai T, Porpiglia E, Lund PJ, Markov G, et al. Glucose Metabolism Drives Histone Acetylation Landscape Transitions that Dictate Muscle Stem Cell Function. Cell Rep. 2019;27: 3939-3955.e6. doi:10.1016/j.celrep.2019.05.092

37. tumanov S, Bulusu V, Gottlieb E, Kamphorst JJ. A rapid method for quantifying free and bound acetate based on alkylation and GC-MS analysis. Cancer Metab. 2016;4: 17. doi:10.1186/s40170-016-0157-5

38. Baeza J, Lawton AJ, Fan J, Smallegan MJ, Lienert I, Gandhi T, et al. Revealing Dynamic Protein Acetylation across Subcellular Compartments. J Proteome Res. 2020;19: 2404–2418. doi:10.1021/acs.jproteome.0c00088

39. Choudhary C, Kumar C, Gnad F, Nielsen ML, Rehman M, Walther TC, et al. Lysine acetylation targets protein complexes and co-regulates major cellular functions. Science. 2009;325: 834–840. doi:10.1126/science.1175371

40. trefely S, Liu J, Huber K, Doan MT, Jiang H, Singh J, et al. Subcellular metabolic pathway kinetics are revealed by correcting for artifactual post harvest metabolism. Mol Metab. 2019;30: 61–71. doi:10.1016/j.molmet.2019.09.004

41. Chen WW, Freinkman E, Wang T, Birsoy K, Sabatini DM. Absolute Quantification of Matrix Metabolites Reveals the Dynamics of Mitochondrial Metabolism. Cell. 2016;166: 1324-1337.e11. doi:10.1016/j.cell.2016.07.040

42. Zhang M, Pan Y, Dorfman RG, Yin Y, Zhou Q, Huang S, et al. Sirtinol promotes PEPCK1 degradation and inhibits gluconeogenesis by inhibiting deacetylase SIRT2. Sci Rep. 2017;7: 7. doi:10.1038/s41598-017-00035-9

43. Ponnusamy M, Zhou X, Yan Y, Tang J, Tolbert E, Zhao TC, et al. Blocking sirtuin 1 and 2 inhibits renal interstitial fibroblast activation and attenuates renal interstitial fibrosis in obstructive nephropathy. J Pharmacol Exp Ther. 2014;350: 243–256. doi:10.1124/jpet.113.212076

44. Bitterman KJ, Anderson RM, Cohen HY, Latorre-Esteves M, Sinclair DA. Inhibition of Silencing and Accelerated Aging by Nicotinamide, a Putative Negative Regulator of Yeast Sir2 and Human SIRT1. Journal of Biological Chemistry. 2002;277: 45099–45107. doi:10.1074/jbc.M205670200

45. Kasamatsu A, Nakao M, Smith BC, Comstock LR, Ono T, Kato J, et al. Hydrolysis of O-acetyl-ADP-ribose isomers by ADP-ribosylhydrolase 3. J Biol Chem. 2011;286: 21110–21117. doi:10.1074/jbc.M111.237636

46. Ono T, Kasamatsu A, Oka S, Moss J. The 39-kDa poly(ADP-ribose) glycohydrolase ARH3 hydrolyzes O-acetyl-ADP-ribose, a product of the Sir2 family of acetyl-histone deacetylases. Proc Natl Acad Sci U S A. 2006;103: 16687–16691. doi:10.1073/pnas.0607911103

47. Khan N, Jeffers M, Kumar S, Hackett C, Boldog F, Khramtsov N, et al. Determination of the class and isoform selectivity of small-molecule histone deacetylase inhibitors. Biochem J. 2008;409: 581–589. doi:10.1042/BJ20070779

48. Yoshida M, Kijima M, Akita M, Beppu T. Potent and specific inhibition of mammalian histone deacetylase both in vivo and in vitro by trichostatin A. J Biol Chem. 1990;265: 17174–17179.

49. Koeller KM, Haggarty SJ, Perkins BD, Leykin I, Wong JC, Kao M-CJ, et al. Chemical genetic modifier screens: small molecule trichostatin suppressors as probes of intracellular histone and tubulin acetylation. Chem Biol. 2003;10: 397–410. doi:10.1016/s1074-5521(03)00093-0

50. Izzo L, Trefely S, Demetriadou C, Drummond J, Mizukami T, Kuprasertkul N, et al. The carnitine shuttle links mitochondrial metabolism to histone acetylation and lipogenesis. bioRxiv. 2022; 2022.09.24.509197. doi:10.1101/2022.09.24.509197

51. Lin CY, Kumar S. Pathway for the Synthesis of Fatty Acids in Mammalian Tissues. Journal of Biological Chemistry. 1972;247: 604–606. doi:10.1016/S0021-9258(19)45745-1

52. Wellen KE, Snyder NW. Should we consider subcellular compartmentalization of metabolites, and if so, how do we measure them? Curr Opin Clin Nutr Metab Care. 2019;22: 347–354. doi:10.1097/MCO.0000000000000580

53. Ryu KW, Nandu T, Kim J, Challa S, DeBerardinis RJ, Kraus WL. Metabolic regulation of transcription through compartmentalized NAD+ biosynthesis. Science. 2018;360: eaan5780. doi:10.1126/science.aan5780

54. Lee DY, Hayes JJ, Pruss D, Wolffe AP. A positive role for histone acetylation in transcription factor access to nucleosomal DNA. Cell. 1993;72: 73–84. doi:10.1016/0092-8674(93)90051-q

55. Kouzarides T. Chromatin modifications and their function. Cell. 2007;128: 693–705. doi:10.1016/j.cell.2007.02.005

56. Lee JV, Berry CT, Kim K, Sen P, Kim T, Carrer A, et al. Acetyl-CoA promotes glioblastoma cell adhesion and migration through Ca ^2+^ –NFAT signaling. Genes & Development. 2018;32: 497–511. doi:10.1101/gad.311027.117

57. trefely S, Ashwell P, Snyder NW. FluxFix: automatic isotopologue normalization for metabolic tracer analysis. BMC Bioinformatics. 2016;17: 485. doi:10.1186/s12859-016-1360-7

58. Snyder NW, Tombline G, Worth AJ, Parry RC, Silvers JA, Gillespie KP, et al. Production of stable isotope-labeled acyl-coenzyme A thioesters by yeast stable isotope labeling by essential nutrients in cell culture. Anal Biochem. 2015;474: 59–65. doi:10.1016/j.ab.2014.12.014

59. Schindelin J, Arganda-Carreras I, Frise E, Kaynig V, Longair M, Pietzsch T, et al. Fiji: an open-source platform for biological-image analysis. Nat Methods. 2012;9: 676–682. doi:10.1038/nmeth.2019

60. Frey AJ, Feldman DR, Trefely S, Worth AJ, Basu SS, Snyder NW. LC-quadrupole/Orbitrap high-resolution mass spectrometry enables stable isotope-resolved simultaneous quantification and (13)C-isotopic labeling of acyl-coenzyme A thioesters. Analytical and bioanalytical chemistry. 2016;408: 3651–8. doi:10.1007/s00216-016-9448-5

