## Supplementary figures and images for "Dynamic protein deacetylation is a limited carbon source for acetyl-CoA-dependent metabolism"

### Supplemental Figure 1

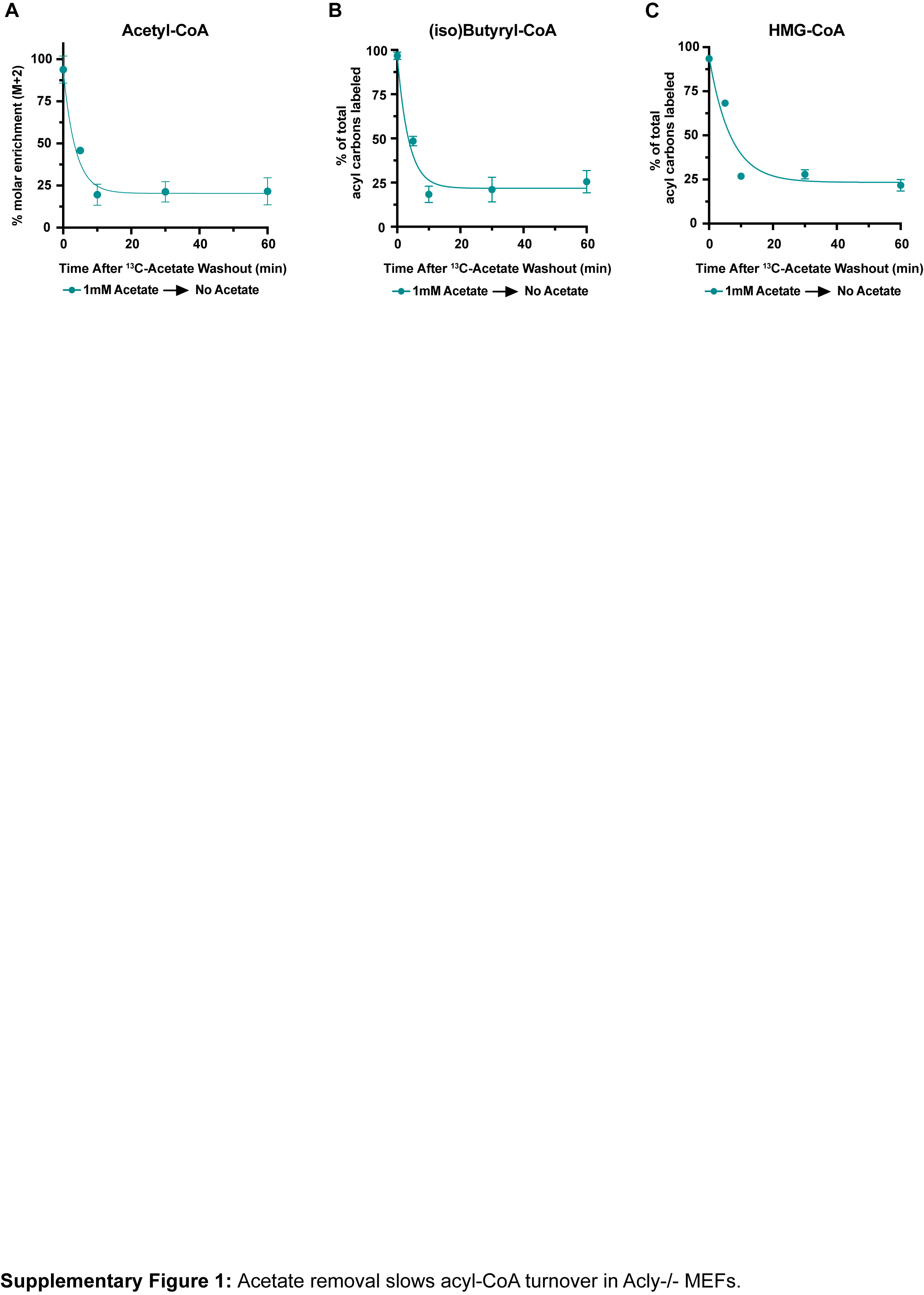

### Supplemental Figure 2

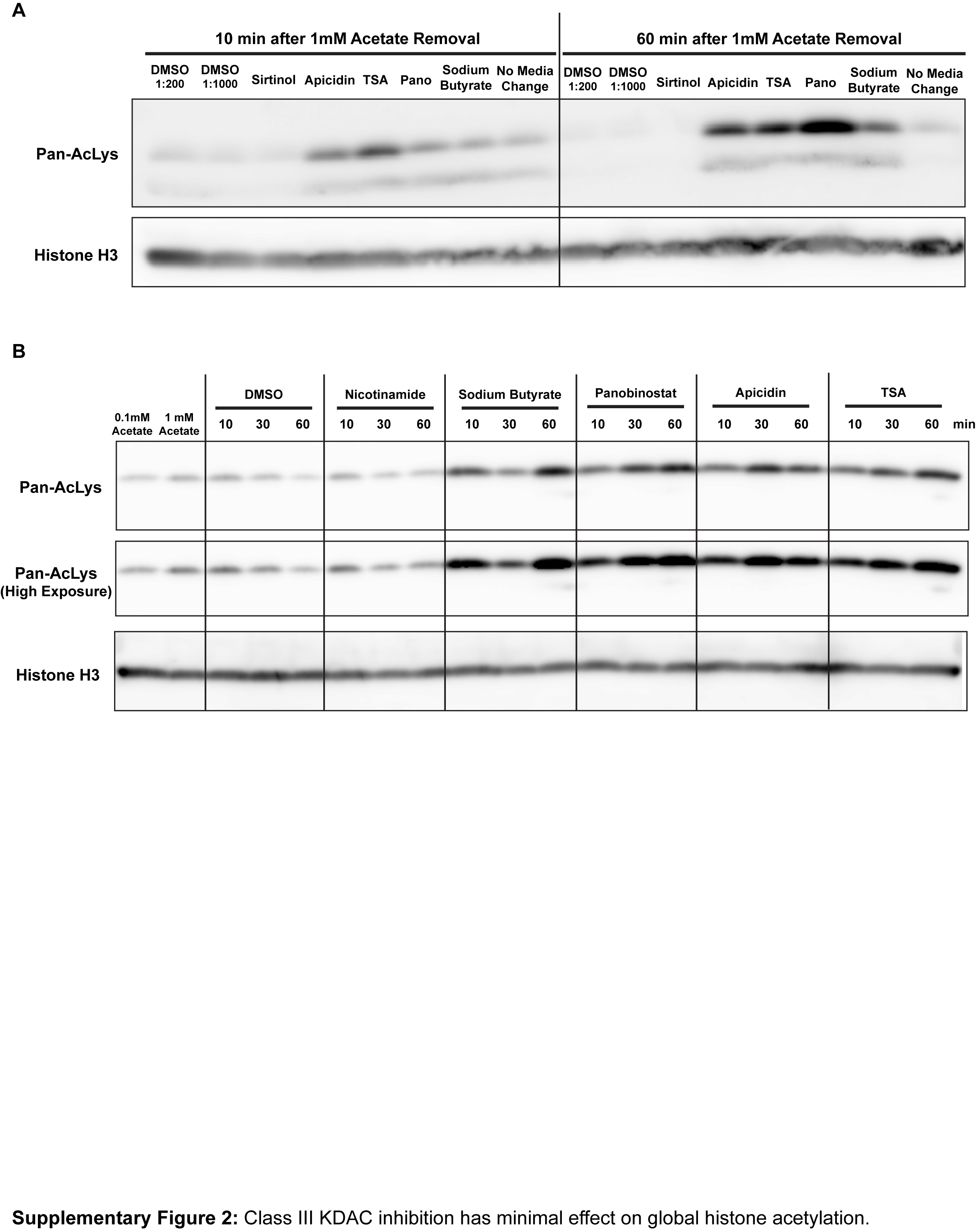

### Supplemental Figure 3

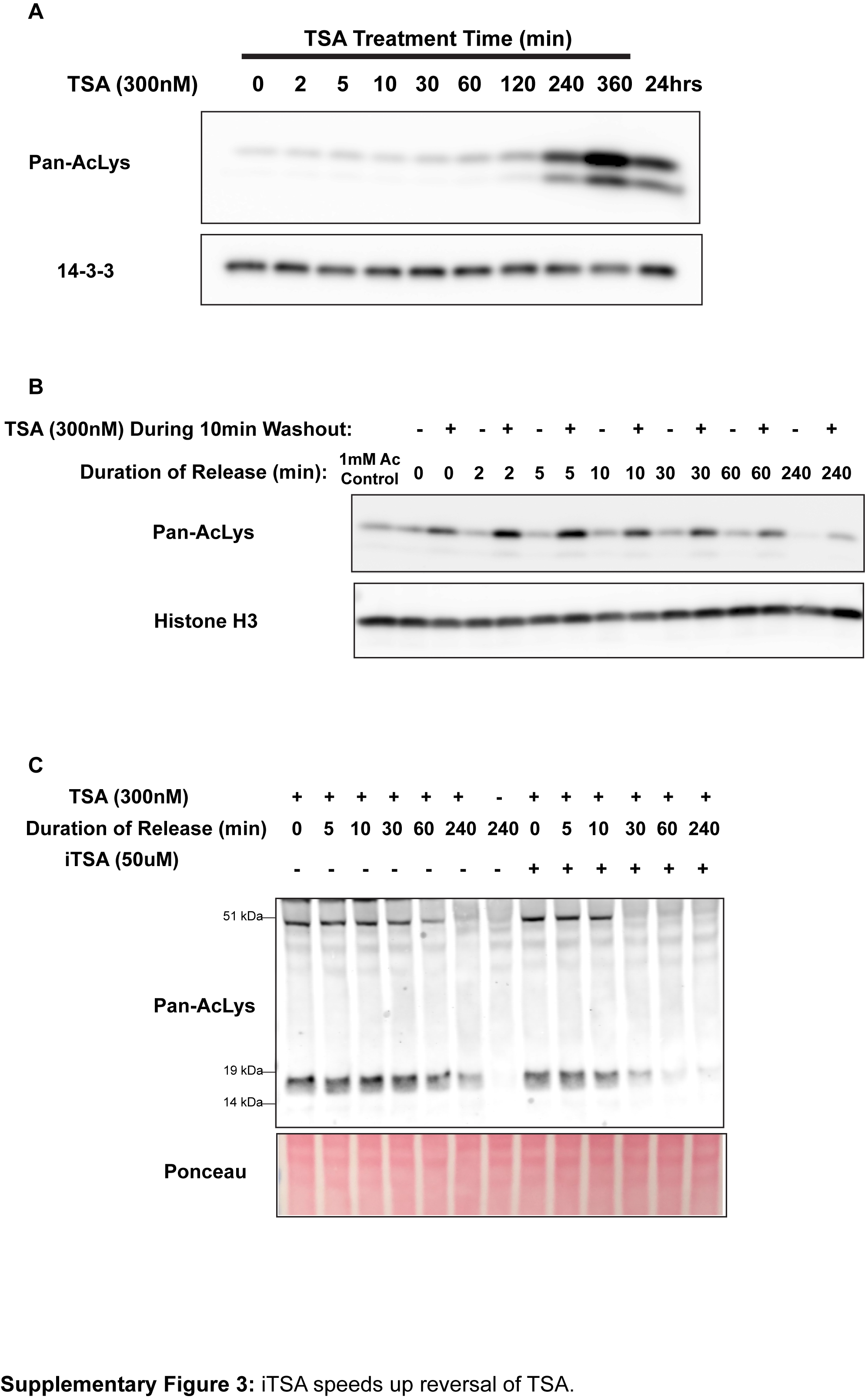

### Supplemental Figure 4

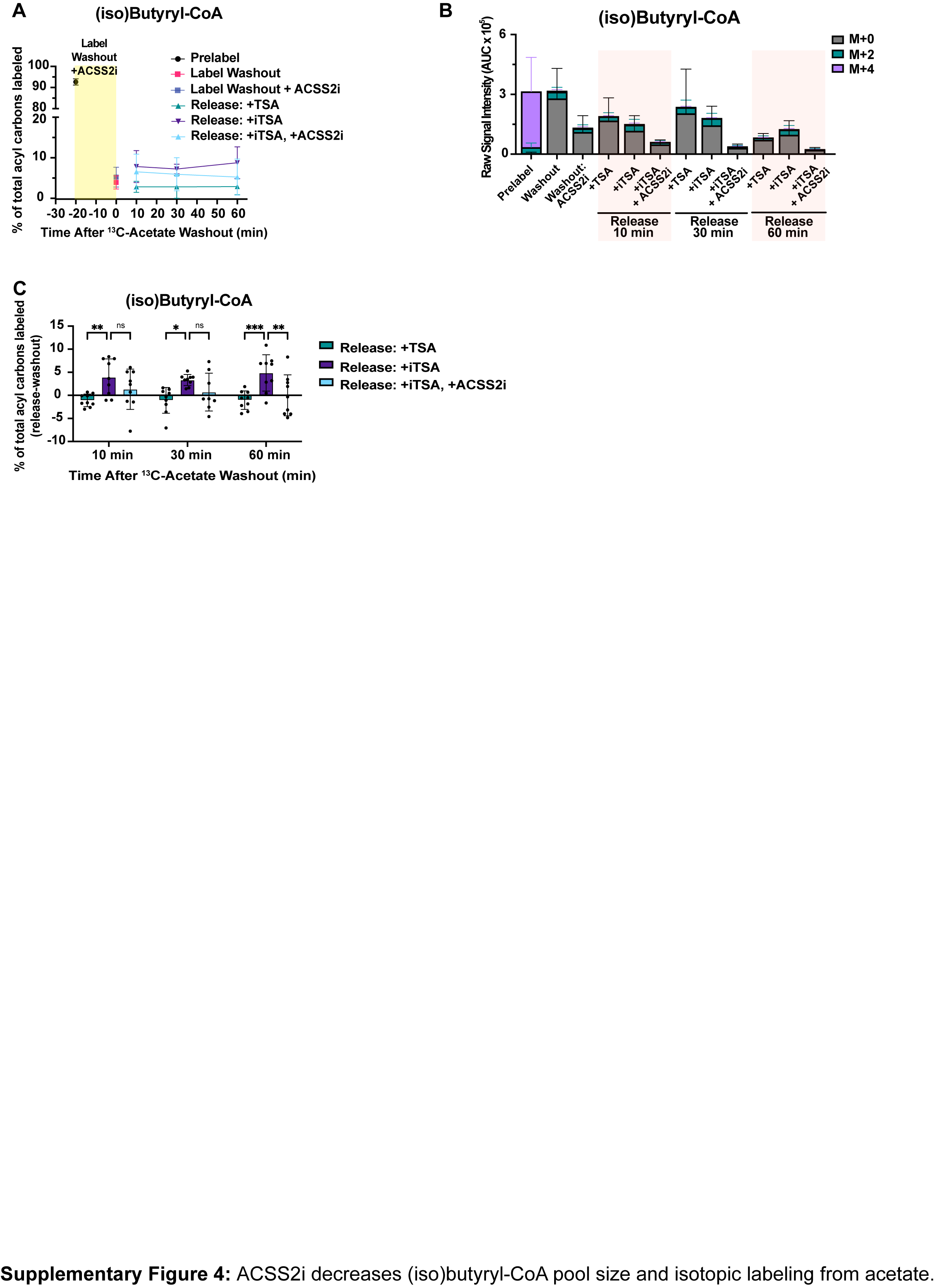
